# Complex plumages spur rapid color diversification in island kingfishers (Aves: Alcedinidae)

**DOI:** 10.1101/2022.09.26.509475

**Authors:** Chad M. Eliason, Jenna M. McCullough, Shannon J. Hackett, Michael J. Andersen

## Abstract

Oceanic islands are cradles for diversity. Differences in predation pressures and lack of competition on islands are thought to drive both phenotypic and species diversification. While most work exploring these patterns has focused on life history, behavioral and morphological traits, many island species are uniquely colorful. Yet, a recent study of island bird coloration found that insular species are duller than continental species. Whether such shifts in color are associated with increased rates of color evolution on islands remains unknown. Here, we incorporate geometric morphometric techniques to study plumage color diversity in a speciose clade of colorful birds that inhabit nearly all areas of the globe—kingfishers (Aves: Alcedinidae). In particular, we test two hypotheses: (i) that plumage complexity enhances interspecific rates of color evolution and (ii) that plumage color diversity is elevated on islands. Our results show that more complex plumages result in more diverse colors among species and plumage color evolves faster on islands. Importantly, we found that insular species did not have more complex plumages than their continental relatives. Thus, complexity may be a key innovation that facilitates response to divergent (or relaxed) selection pressures on islands. Lack of support for competition driving rates of evolution along different color axes hints at an allopatric model of color evolution in which species adapt to local conditions on different islands. This work demonstrates how a truly multivariate treatment of color data can reveal evolutionary patterns that might otherwise go unnoticed.

## Introduction

Islands are natural laboratories for studying evolution in action (Losos and Ricklefs, 2009). Since they often lack natural predators and competitors due to their geographic isolation, island systems enable colonizing species to occupy new niches that are not available on the mainland. Adaptation to these novel niches is predicted to lead to both more species and changes in phenotypes (Schluter, 2000). The “island rule” predicts that large bodied colonists evolve to be smaller on islands than their mainland congeners (dwarfism) and vice versa for smaller-bodied colonists to evolve to be large (gigantism). While less commonly assessed, these shifts in phenotype when moving to islands are likely accompanied by bursts in phenotypic evolutionary rate (Millien, 2006; Thomas et al., 2009; Woods et al., 2020). Another common feature of evolution on islands is the tendency for convergent evolution “island syndrome”), for example, in life history traits (Covas, 2012; Losos et al., 1998; Novosolov et al., 2013), behavior (Buglione et al., 2019; Roff, 1994), and morphology (Clegg and Owens, 2002; Wright et al., 2016).

Lower intensity of sexual selection and differences in predation pressure or resource availability on islands set the stage for shifting targets of selection on intraspecific signals within island systems. Compared to life history and morphological traits, intraspecific signals used in mating and social contexts are not commonly explored in the context of island evolution. Previous work has shown increased color polymorphism in island snails (Bellido et al., 2002; Ożgo, 2011) and lizards (Corl et al., 2010). Within birds, island species tend to be less sexually dimorphic and have simpler songs (Price, 2008). Island birds have also been shown to have drabber colors and simpler plumage patterns (Bliard et al., 2020; Doutrelant et al., 2016). Yet, we lack a detailed understanding of color evolution within, rather than between, island and mainland clades. Moreover, color patterns can themselves be influenced by multiple selective factors (Cuthill et al., 2017). Since selection can only act on existing variability, such as distinct patches within a plumage, ancestrally shared developmental bases of plumage patterns might act as a brake on color evolution (Price and Pavelka, 1996). For a uniformly colored species, selection on color would presumably cause the whole plumage to change in tandem. By contrast, if a species is non-uniformly colored (i.e., patchy, and therefore more complex), selection can act on different aspects of coloration (Brooks and Couldridge, 1999). On a macroevolutionary scale, we would predict greater color divergence in a clade with an ancestrally complex plumage pattern because there is more heritable color variation (i.e., among patches) upon which selection can act. Plumage complexity, therefore, could be a key innovation driving rates of color evolution in birds.

Generally, birds produce color by two different mechanisms—light absorption by pigments and light scattering by feather nanostructures (Shawkey and D’Alba, 2017)—and these have been shown to have different effects on plumage evolutionary rates. Melanin, the most prevalent pigment in avian plumages (Mcgraw, 2006), produces the black, brown, gray, and rufous colors. The production of highly complex barring and mottled plumages stems from melanin within feathers. These pigments have an important role in the ecology of birds, as melanin-based pigments increase the durability of feathers (Bonser, 1995) and aid as a protectant against degradation (Goldstein et al., 2004). Birds also incorporate carotenoid-based color, which is exclusively derived from a bird’s diet (Olson and Owens, 1998), to produce red, orange, and yellow colored plumages. Since carotenoid-based color is so closely linked to an individuals’ health and diet, these plumage colors are considered an honesty-reinforcing, “true” signal of an individual’s fitness (Lank and Hill, 2004; Simons et al., 2014; von Schantz et al., 1999). Whereas melanin and carotenoid-based coloration are produced by chemical pigments and absorb light waves, structural colors are produced by the physical interaction of light waves and nanometer-scale variations in the feather integument and the scattering of light (Prum, 2006). Within birds, structural colors produce a wide array of color, including blue-green colors, glossy blacks, as well as iridescence. Because structural colors are more evolutionarily labile than pigment-based colors, they have faster evolutionary rates and are considered key innovations (Maia et al., 2013b).

Two hallmarks of kingfishers (Aves: Alcedinidae) are their complex plumage patterns (Eliason et al., 2019) and their island distributions (Andersen et al., 2018; McCullough et al., 2019). Kingfishers encompass a wide variety of colors—from the aquamarine-colored back of the common kingfisher (*Alcedo atthis*), to the brilliant silver back of the southern silvery-kingfisher (*Ceyx argentatus*), or the purple rump of the ultramarine kingfisher (*Todiramphus leucopygius*). They also run the gamut of plumage complexity, including intricate scalloping of the spotted kingfisher (*Actenoides lindsayi*) and the contrasting hues of the black-backed dwarf-kingfisher (*Ceyx erithaca*). The family is widely distributed across the globe but their center of diversity is Indomalaya, including island clades in Wallacea and Melanesia recently having been highlighted for their high diversification rates (Andersen et al., 2018). These same island clades, specifically within the woodland kingfisher genus *Todiramphus*, also have elevated color disparity between closely related species (Eliason et al., 2019). Smaller population sizes, isolation, and genetic drift could promote color divergence within island kingfishers, contributing to high rates of color evolution and disparity. Kingfishers could be an ideal system to investigate interactions between key innovations (complex plumages) and color diversification in island systems.

In this study, we implement geometric morphometric techniques to investigate complex plumage pattern evolution across kingfishers. We tested whether plumage complexity increases interspecific rates of color and predict that lineages with a more ancestrally complex plumage pattern would have higher overall rates of plumage evolution. In addition, we tested whether island systems, which experience less predation and relaxed sexual selection pressures, would have higher plumage diversity than closely related species on continental systems. We predicted that island colonization would result in faster rates of color evolution compared to mainland species. Our study of the interplay between the arrangement of color patches (a potential key innovation) and geography sheds light more broadly on the role of spatial opportunity in phenotypic evolution.

## Results

### Holistic assessment of plumage color pattern variation

Plumage coloration is highly multivariate, varying both within feathers, among feather regions on a bird, between sexes, and among species. To visualize major axes of variation in overall plumage color patterning, we used a phylogenetic principal components analysis (pPCA), with per-patch color coordinates as variables (N = 66). We then plotted the first two pPC scores that together accounted for >50% of color variation in the clade (Figure 1; see Figure S1 for ordinary PCA plot).

**Figure 1.**
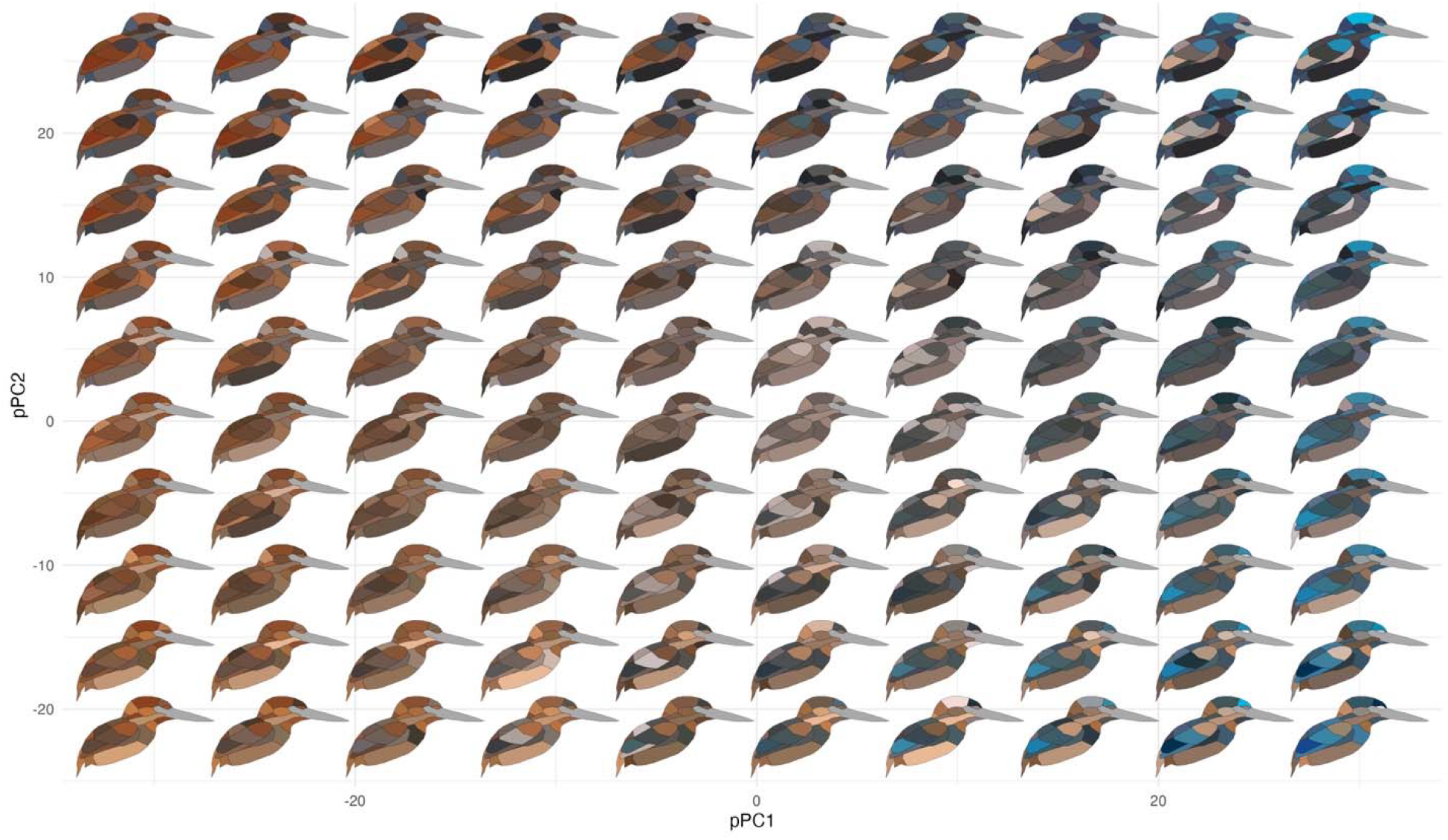
Color pattern morphospace of kingfishers. Bird images show depictions of color in a human visual system based on spectral measurements over a grid of phylogenetic principal components analysis (pPCA) coordinates. Axes shown are pPC axes 1 and 2, together accounting for >50% of plumage color variation in the clade.

To understand how color varies across these different levels, we performed a taxonomic ANOVA (Derrickson and Ricklefs, 1988). Briefly, we fit a linear mixed model in MCMCglmm using colorspace XYZ coordinates as a multivariate response, with random effects for plumage patch, sex, and species. We then estimated the proportion of variation explained at each level of organization (see Dryad for R code). Partitioning of variance analysis revealed three distinct modes of color variation within kingfishers: (i) clades with complex color patterns that partition color variance more among patches than among species or individuals (e.g., *Dacelo*, Alcedininae, Cerylininae), (ii) clades that vary primarily among individuals/sexes (e.g., *Tanysiptera*), and (iii) clades that vary mainly among species (e.g., *Todiramphus, Corythornis*; Figure 2B).

**Figure 2.**
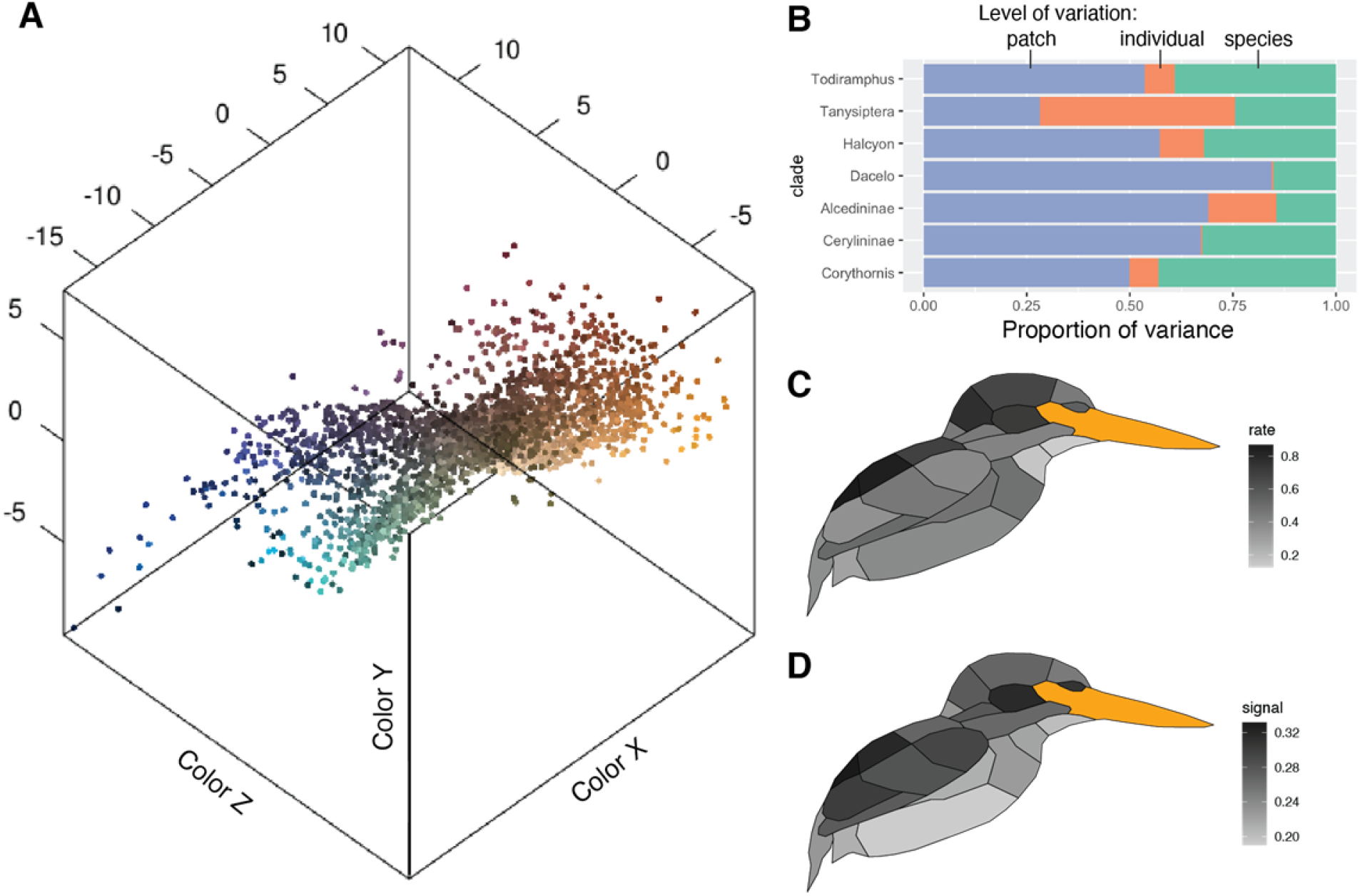
Perceptually uniform colorspace and color variation in kingfishers. (A) Color data, with points being the average of three plumage patch measurements for each individual (N = 3101). Colors are estimated from a human visual system using spec2rgb in pavo (Maia et al., 2013a). Distance between patches are proportional to the just noticeable distances (JNDs), assuming a UV-sensitive visual system (Parrish et al., 1984). (B) Proportional color variance among patches in an individual (orange), among individuals in a species (green), and among species in a clade (violet). (C) Distribution of multivariate evolutionary rates and (D) phylogenetic signal of color evolution across the body (darker colors indicate higher values).

We next compared phylogenetic signal and rates of color evolution within each individual plumage patch using distance-based comparative methods (Adams, 2014; Denton and Adams, 2015). This analysis showed that rates of color evolution are unevenly distributed across the body, with dorsal regions evolving faster than ventral ones (Figure 2C). This differs from several previous studies illustrating rapid rates of ventral plumage evolution in tanagers (Shultz and Burns, 2017), manakins (Doucet et al., 2007), fairy-wrens (Friedman and Remeš, 2015), and antbirds (Marcondes and Brumfield, 2019). Thus, dorsal plumage patches may be under stronger sexual selection in kingfishers, as rapid rates of display trait evolution are thought to be associated with more intense sexual selection (Irwin et al., 2008; Seddon et al., 2013). Rump and cheek patches showed the highest levels of phylogenetic signal, suggesting that these patches are more diagnostic of species than crown or wing plumage (Figure 2D).

### A new approach for comparing plumage complexity to rates of color evolution

To test our hypothesis that plumage complexity facilitates interspecific color divergence, we required ancestral states of color complexity to work with existing methods that calculate rates for each node in a phylogeny (Revell, 2011). To do so, we used a phylogeny by McCullough et al. (2019), which incorporated thousands of ultraconserved elements (Faircloth et al., 2012) for a fully sampled, time-calibrated phylogeny of the order Coraciiformes. For each “trait” (i.e., unique combination of coordinate XYZ axis and patch), we reconstructed ancestral states using the ace function in the R package ape (Paradis et al., 2004). We then calculated plumage complexity at each node/tip in four ways: as the mean pairwise distance among all patches in XYZ space (at tips and nodes based on ancestral XYZ values); as the color volume enclosing all points for a species, and as the number of perceptually distinguishable (just noticeable difference, JND > 1) and contiguous color regions on the body for both a folded-wing and spread-wing plumage configuration. The latter two metrics are similar to a recent method (Eliason et al., 2019) of calculating color complexity of plumages as the number of contiguous body regions sharing the same color mechanism (e.g., melanin-based or structural coloration), but they are based on continuous reflectance values instead of discrete color data (i.e., presence or absence of a given color mechanism). Higher differences between adjacent patches yielded higher plumage complexity scores.

Ancestral estimates of color complexity were correlated among our four different complexity metrics (Figure S2). To obtain per-node multivariate evolutionary rates, we computed multivariate independent contrasts following McPeek et al. (2008) with an extension for avian tetrahedral colorspace (see Dryad for R code). At each node, we calculated the perceptual color distance between both daughter lineages in avian colorspace using the coldist function in pavo (Maia et al., 2013a). We finally divided this divergence by the total branch lengths (time) between each daughter lineage (Felsenstein, 1985). Given these two sets of data (ancestral evolutionary rates and estimates of plumage complexity), we can now test for a relationship between them by comparing the observed correlation to that obtained using simulation data (see Supplemental Methods).

### Species with complex plumages have higher rates of plumage color evolution

Plumage complexity and interspecific differences in coloration are typically thought of as distinct axes of color diversity. Yet, species that have evolved several patches have more “degrees of freedom” to vary. Here, we attempt to link complexity with interspecific variation (i.e., rates) using a novel approach with multivariate data. Combining ancestral estimates of color complexity and overall plumage divergence, we found that rates of color evolution were higher in species with more complex plumages (Figure 3C, Table S2). This finding is consistent with the idea of multifarious selection providing more axes for ecological or phenotypic divergence among species, and can eventually lead to speciation (Nosil et al., 2009). However, recent work in wolf spiders has revealed that signal complexity per se can be a direct target of sexual selection (Choi et al., 2022). Another possibility in kingfishers is that body size is driving the evolution of plumage complexity, as signal complexity has been shown to decrease with body size in iguanian lizards (Ord and Blumstein, 2002). Interestingly, the kingfisher species with the most complex plumages are also among the smallest birds in the family, such as the indigo-banded kingfisher (*Ceyx cyanopectus*) and Southern silvery-kingfisher (*Ceyx argentatus*, Figure 3). Alternatively, if species on islands have more complex plumages, then islands might indirectly drive color divergence. However, a statistical comparison of plumage complexity metrics between islands and mainland taxa showed no support for this idea (Table S3, Figure S3). This suggests that island-dwelling and plumage complexity may be independent drivers of color variation in the group.

**Figure 3.**
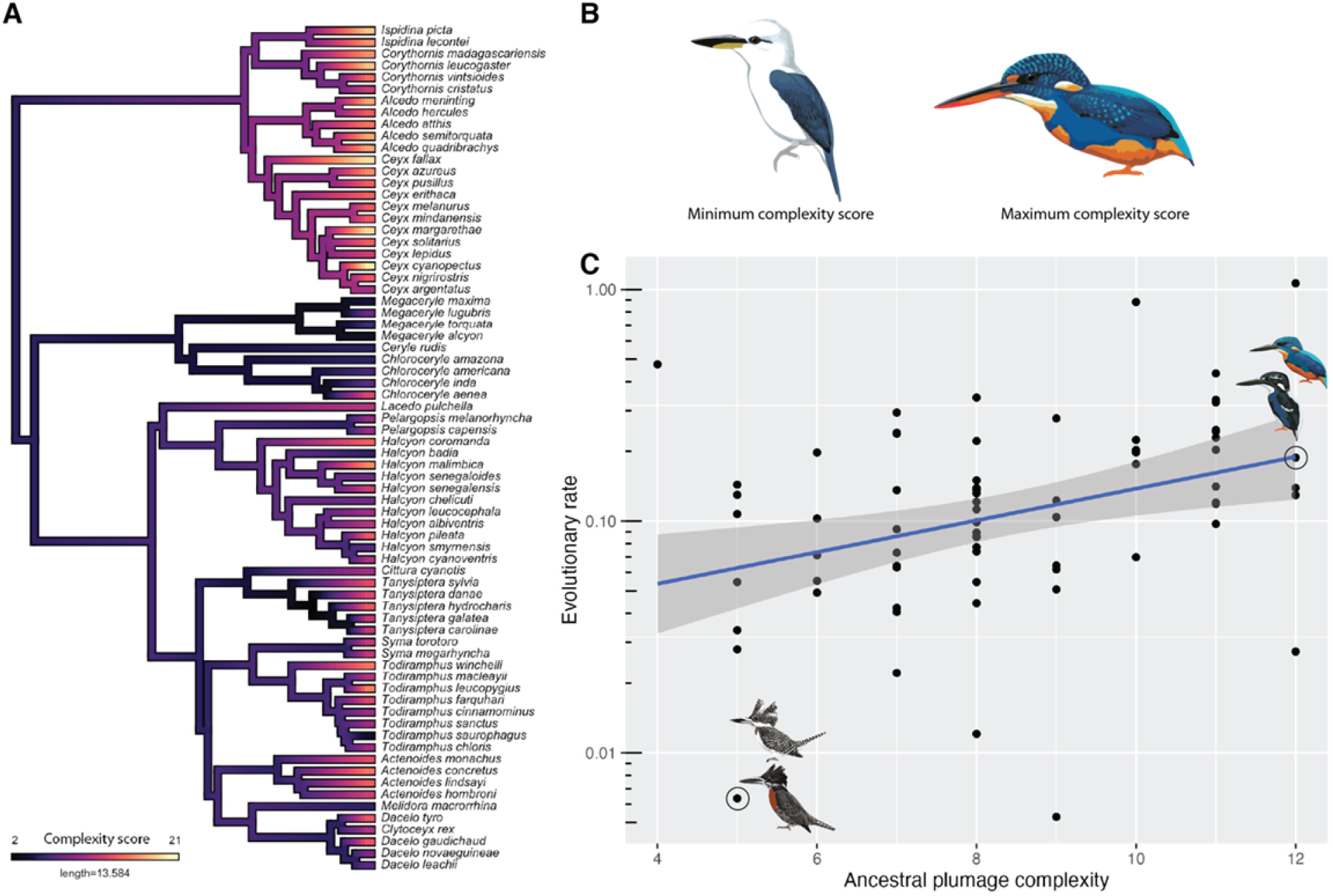
Species with complex plumages evolve color variation at a faster rate. (A) Evolution of plumage color complexity (results shown for spread-wing configuration). Complexity calculated as the number of distinct contiguous color patches in a species (N = 22 patches). (B) Species with minimum (beach kingfisher [*Todiramphus saurophagus*]) and maximum (indigo-banded kingfisher [*Ceyx cyanopectus*]) values of plumage complexity. (C) Relationship between overall color divergence between sister lineages and spread-wing plumage complexity at that node (Spearman’s ρ = 0.37, P < 0.01). Two species pairs are shown to illustrate weak color divergence in simple plumage patterns (left) and strong divergence in species with complex plumage patterns (right). Illustrations created by Jenna McCullough.

### Island kingfishers have higher rates of color evolution

To test for an “island effect” on rates of color evolution, we treated individual patches as geometric morphometric landmarks and compared multivariate evolutionary rates between insular and continental species using the compare.evol.rates function (Denton and Adams, 2015) in geomorph (Adams and Otárola-Castillo, 2013). Models were fit using the ML estimate of the branch length transformation parameter Pagel’s λ. To further test the influence of phylogenetic signal on evolutionary rate comparisons, we transformed branch lengths over a range of values for Pagel’s λ, following Adams (2014). Briefly, we transformed branches with λ ranging from 0–1 (in 20 equidistant steps) and then refit evolutionary rate models. Since biased sampling can also influence evolutionary rates (e.g., if our sampling captures extremes of population color distributions), we further accounted for measurement error in our analyses by estimating intraspecific variation in the collared kingfisher (*Todiramphus chloris*). We measured 55 individuals from 13 subspecies and used these measurements to calculate standard errors of XYZ coordinates in colorspace for each plumage patch. Following Denton and Adams (2015), we created pseudosamples of specimens by randomly sampling values within the range of species mean ± standard error for *Todiramphus chloris*. We repeated this 1000 times, each time calculating species means and re-running rate tests. We then compared this distribution of rates for each regime (islands, mainland) to that estimated from the original species means. We found that species distributed on islands are evolving color faster than continental species (P = 0.037, σ^2^_cont_ = 0.133, σ^2^_island_ = 0.216; Figure 4). This result was robust to measurement error (Figure 4, inset) and variable phylogenetic signal (Figure S4).

**Figure 4.**
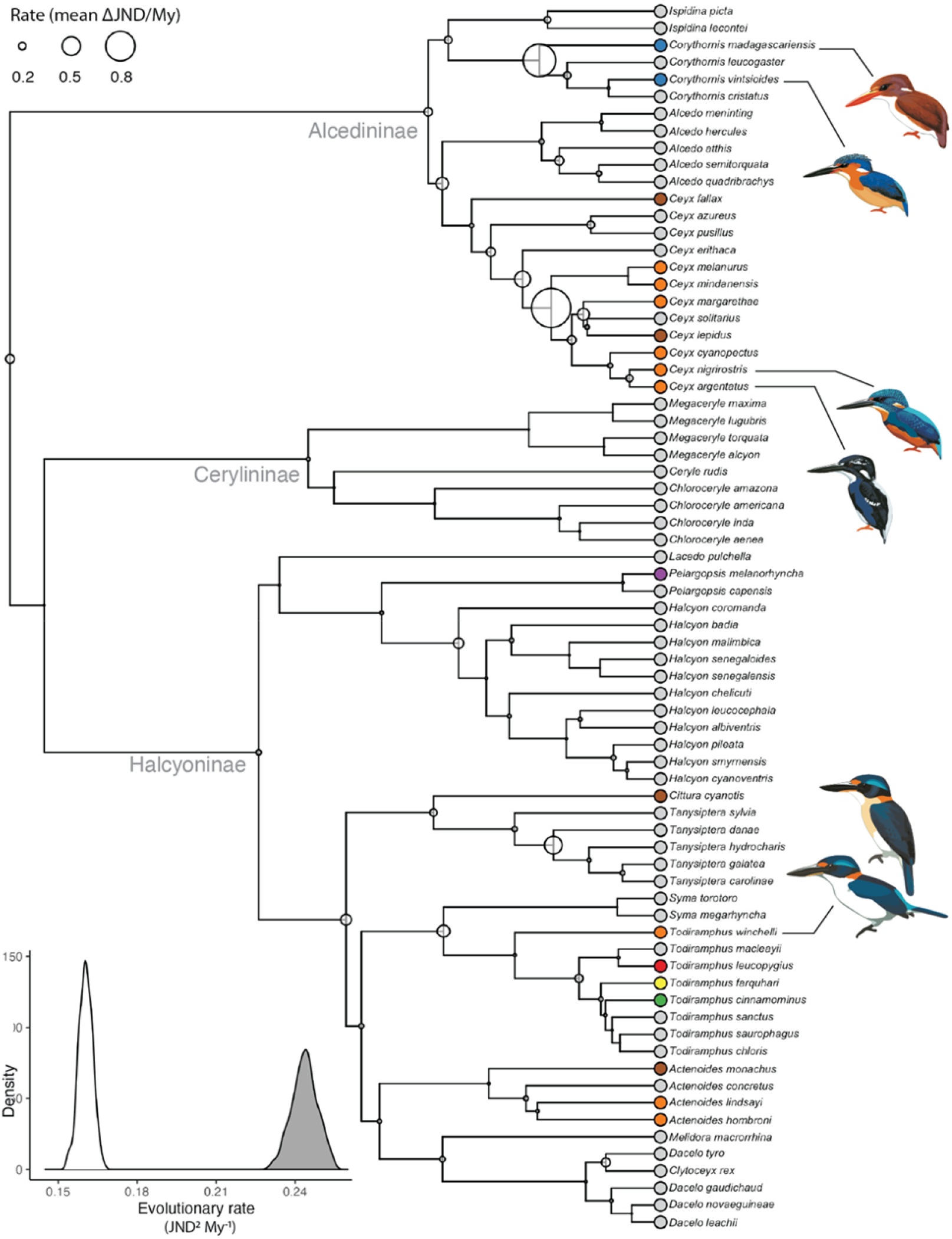
Rapid rates of plumage color evolution in island kingfishers. Phylogeny of kingfishers (N = 72 species) with circles at nodes proportional to mean interspecific patch color divergence among species. Tip colors correspond to different islands (see Fig. 3 legend), with continental species in gray. Inset shows effect of measurement error on divergence in rates of evolution between continental (white) and island species (grey) The relationship is significant (rate ratio = 1.62, P = 0.037). Illustrations created by Jenna McCullough.

To test the possibility that rapid color divergence on islands is the result of reproductive character displacement within islands (e.g., see Losos and Ricklefs, 2009), we fit and compared models of trait evolution that do and do not account for competition among species. First, we ran a phylogenetic principal components analysis (pPCA) and used the first 5 pPC components that together accounted for >95% of color variation in comparative analyses. Next, for each pPC score and two geographic species subsets (islands only, mainland only), we fit five models of trait evolution: a Brownian motion (BM) model, an Ornstein-Uhlenbeck (OU) model, a matching competition (MC) model, a density-dependent model in which trait divergence (rate) is a linear function of the number of sympatric species (DD_lin_), and a density-dependent model in which trait divergence is an exponential function of the number of sympatric species (DD_exp_). For the full species data set, we fit these same five models in addition to a sixth model: a multi-regime BM model in which rates and means are allowed to vary between island and mainland taxa (BMS). We found no support for interspecific competition driving color divergence among species, as the OU model was the best fitting model for all color pPC axes (Figure 5C). The pattern was slightly different within islands, although AICc weights were ∼0.6 suggesting weak support for competition driving color evolution on islands (Figure 5B). Taken together, this suggests that intraspecific competition may instead be driving color in island kingfishers.

**Figure 5.**
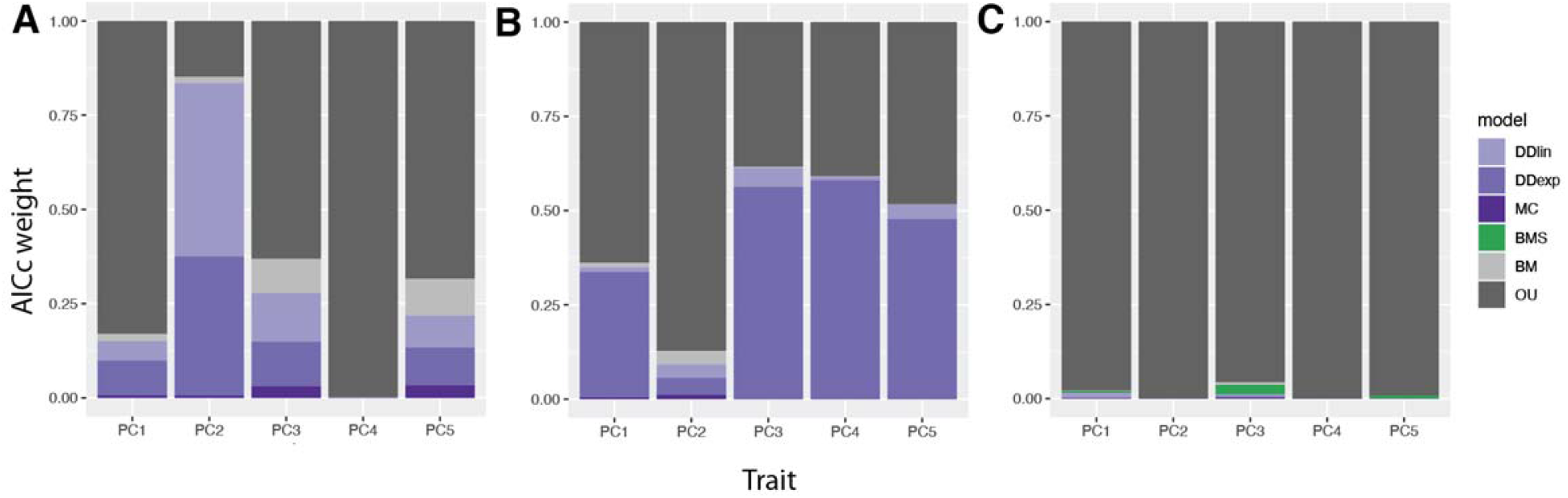
Interspecific competition is not a major driver of color divergence on islands. Relative support for different models of trait evolution. Continental species only (A), island species only (B), and all lineages (C). There is overall ambiguous support (AICc weights ∼0.6) for competition driving color evolution (e.g., DD_lin_, DD_exp_, and MC models). Note: the BMS model is only fit using the “all lineages” data set because this model depends on a mix of both island and continental species to compare rates of evolution.

### Ecological drivers of color evolution

Predictable features of islands, such as island area, age, or degree of isolation, may drive color evolution in similar ways across island systems. For example, island systems generally have fewer predators and novel selective regimes compared to mainland systems. To test whether islands act as distinct selective regimes and cause convergent evolution in plumage patterns, we used the *convrat* function in the convevol R package (Stayton, 2015). We focused on the C1 convergence metric, calculated as 1 - (D_tip_/D_max_), where D_tip_ is the multivariate distance between two lineages and D_max_ is the maximum distance that those two lineages have obtained at any point since their divergence from a common ancestor. We found no support for the prediction that island species, regardless of clade, should converge on the same plumage color patterns (P = 0.08; see Figure 6). Plumage brightness was also not significantly different between mainland and island species (Figure S5). This is distinct from Doutrelant et al. (2016), who showed predictable trends toward drabber and less colorful plumages on islands.

**Figure 6.**
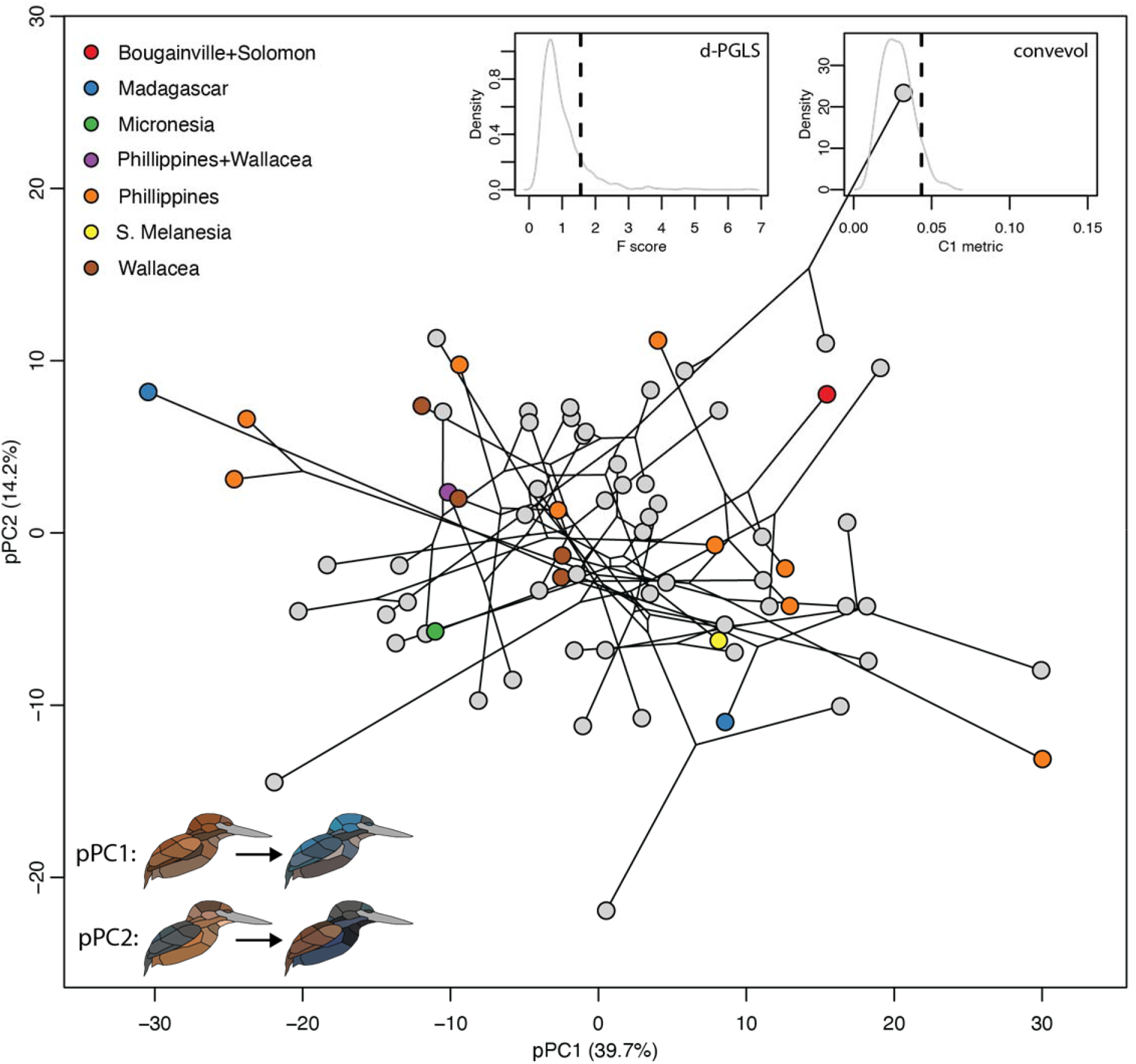
No convergence of color patterns on islands. Phylogenetic principal components analysis (pPCA) plot with points colored by continental (gray) and island species (see legend). Plots in the upper right show no support for global (as measured by the F statistic from a d-PGLS analysis; F = 1.56, P = 0.13) or local (i.e., clade-specific) convergent evolution of color on islands (as measured by the C1 convergence metric; C1 = 0.04, P = 0.08). See inset and Fig. 1 for interpretation of pPC values.

To further understand the ecological and anatomical drivers of color variation, we used Bayesian phylogenetic mixed models (BPMMs). Since structural colors are responsible for most blue and violet colors in nature, we controlled for color-producing mechanism by including it as a covariate in these models. As expected, color-producing mechanism (structural or pigment-based) was a strong predictor of color, with structural colors producing a gamut of turquoise, violet, and blue hues (P < 0.01; Figure 7C). We also found significant sexual dichromatism in color patterns, with males being more blue (P < 0.01; Figure 7A). Color divergence between continental and island species was not significant (P = 0.21; Figure 7B). This echoes the finding of Doutrelant et al. (2016) in a worldwide study, and is also consistent with our finding of no convergence in island color patterns (Figure 6).

**Figure 7.**
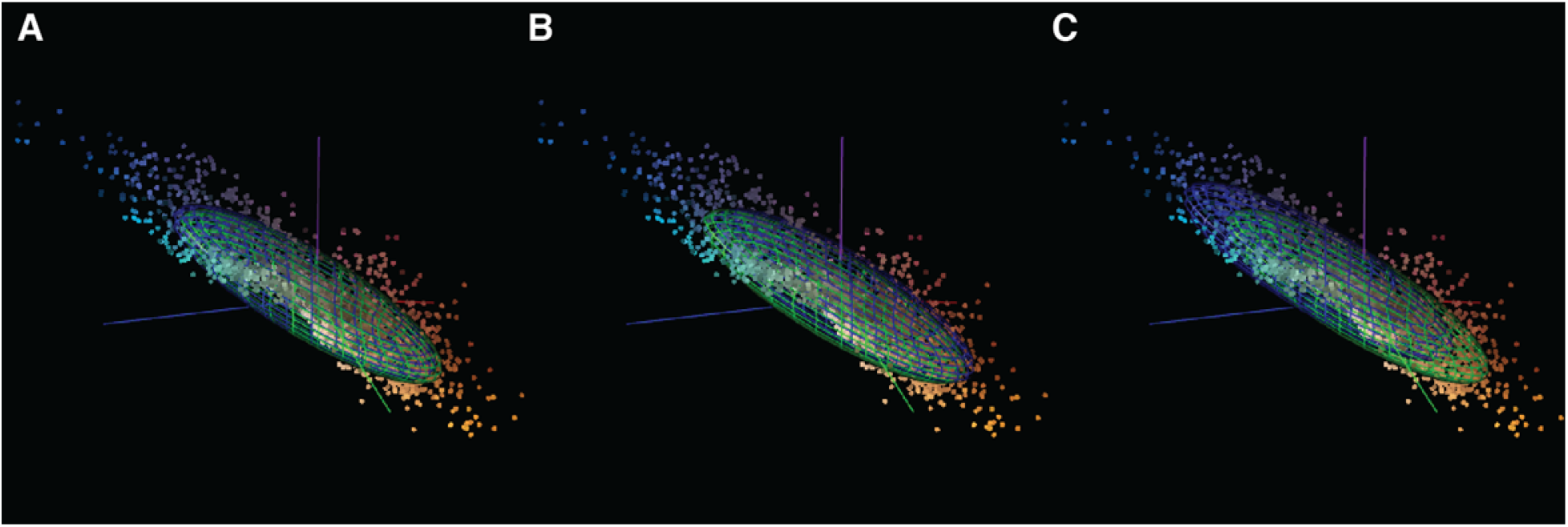
Ecological drivers of color evolution in kingfishers. Results of Bayesian phylogenetic mixed models (BPMM). Panels show effects of sex (A), island-dwelling (B), and color mechanism (C) on plumage color divergence among species. Males (blue ellipsoid) have significantly shorter wavelength colors than females (green ellipsoid; P = 0.01). Species on islands (blue ellipsoid) are not significantly different from continental taxa (green ellipsoid; P = 0.21). As expected, patches with a structural color mechanisms (blue ellipsoid) are significantly more blue than patches with pigment-based color mechanisms (green ellipsoid; P < 0.01).

## Discussion

We lack a cohesive understanding of how plumage color patterns evolve in birds. This study is the first attempt to link color variation among patches (intraspecific) to color variation among species. We find support for higher rates of plumage evolution for clades with more complex plumages, supporting the theory that plumage complexity, rather than uniformity, provides more phenotypic traits for natural selection to act upon. In addition, we find that island lineages have faster rates of plumage evolution than continental lineages.

Colonization of novel geographic areas can promote either shifts in mean phenotype or changes in rates of phenotypic evolution (Collar et al., 2009). Changes in rates associated with island colonization have been described in lizards (Pinto et al., 2008) and mammals (Millien, 2006). Pinto et al. (2008) found that rates of morphological evolution were not elevated in Caribbean anoles compared to mainland species, but they did show differences in morphospace (i.e., convergence). In birds, Doutrelant et al. (2016) measured coloration in 4448 patches of 232 species (including eight kingfisher species) and found that island-dwelling species have duller colors, lower plumage color diversity, and fewer color patches relative to mainland species. This is different from the results of our study, as we show no difference in average plumage brightness (Figure S5) or plumage complexity between mainland and island species (Figure S4, Table S3). Rather, it is the rates of evolution that increase once kingfisher species colonize islands (Figure 4). This suggests decoupling between rapid color evolution on islands and evolutionary changes in plumage complexity, and is consistent with previous work showing elevated rates of morphological evolution being independent of the acquisition of a key innovation, such as toepads of geckos (Garcia-Porta and Ord, 2013). However, this does not answer the question of why plumage color among island lineages would differ more than among mainland lineages.

Colors might evolve for conspicuousness and efficiency in mating displays or species recognition (Andersson, 1994; Doucet et al., 2007). The species-recognition hypothesis predicts reduced signal distinctiveness on islands (West-Eberhard, 1983). This is because of the lack of potential competitors and conspecifics on islands that would otherwise put selective pressures on color patterns, for example through reproductive character displacement (Drury et al., 2018). However, we find the opposite, with more distinct and colorful plumage color signals evolving on islands compared to continental kingfisher taxa (Figure 4). One potential explanation for this pattern is that competition between individuals of the same species is driving color diversity in island kingfishers. That we do not find any support for interspecific competition driving color evolution in kingfishers (Figure 5B) further hints at a role of intraspecific competition driving color in island kingfishers. Other examples of this pattern include island *Anolis* (Gorman, 1968) and *Tropidurus* lizards that show high diversity in dewlap displays (Carpenter, 1966). In birds, parrots are known to have distinctive allopatric and island populations (Forshaw, 1973). At the family level, there are many instances of sympatry on a single island as well as those that are allopatric and endemic to single islands. This pattern is magnified in the genus *Todiramphus*, in which there are many instances of allopatric, island-endemic taxa as well as sympatric cases in which upwards of five *Todiramphus* taxa occur on the same island. In addition, this genus harbors several “superspecies”—a monophyletic group of allopatric taxa too morphologically distinct to be described as a single species (Mayr, 1963)–and are an ideal system in which to test ideas about the role of ecology and constraints driving color diversity within and between islands.

Studying color pattern evolution has been historically difficult, due, in part, to an inability of humans to perceive UV color (Eaton, 2005) and difficulties with measuring and analyzing complex color patterns (Mason and Bowie, 2020). Recent work has showed that changes in plumage complexity are associated with shifts in light environment (Maia et al., 2016; Shultz and Burns, 2013). Our results provide a contrast to these studies in looking at a potential developmental constraint–how plumage patches are arranged on the body (Price and Pavelka, 1996)–and its influence on evolutionary trends of color divergence. Our approach further differs from previous work in that, instead of treating complexity as a univariate trait (Ligon et al., 2018; Maia et al., 2016; Shultz and Burns, 2013), we treat individual plumage patches separately, reconstruct their evolution, and then calculate node-wise plumage complexity scores rather than just at the tips of the tree. With this approach, we recover weaker correlations between color evolutionary rates and ancestral states of plumage complexity when using node complexity scores compared to tip-only complexity scores. The null distribution was also wider for the simulated node data set (Figure S6). This suggests that tip-only complexity data could produce inflated estimates for the relationship between evolutionary rates and plumage complexity than what is actually present in the data (i.e., Type II statistical error). Our node approach is similar to the idea of reconstructing climatic ranges for ancestral species by first reconstructing the ancestral states of individual climate variables (Vieites et al., 2009). One caveat with our approach is that it does not take into account color patterning within feathers. For example, the species with the least complex plumage according to the mean interpatch color distance metric is the crested kingfisher (*Megaceryle lugubris*; see Figure 3C), despite its strong barring/spotting on individual feathers. We hope that researchers will consider the morphometrics approach we take here, as well as assess its potential strengths and weaknesses, in future studies on the evolution of complex color patterns in nature.

In this study we have collected a large amount of spectral data (9303 measurements of 142 individuals in 72 species) in a diverse family of birds notable for their complex plumages and rapid diversification on islands. The major findings of our study are that (i) complex plumage patterns evolve novel colors at a faster rate than simpler uniform plumages and (ii) colonization of islands, independent of plumage complexity, results in further divergence of coloration among species. More broadly, these results highlight the interplay between a potential key innovation (plumage complexity) and geographical opportunity for allopatric speciation in birds. Further research is needed to test whether complex plumages are more common in clades that are speciating rapidly and if complexity is a direct target of sexual selection.

## Materials and methods

### Measuring feather color

Using a UV-Vis spectrophotometer (Ocean Optics) operating in bird-visible wavelengths (300– 700 nm), we measured reflectance spectra in triplicate for 22 patches in 72 species, including both males and females, from museum specimens. In total, we obtained 9303 spectra for 142 individuals (available on Dryad). We then averaged three spectra for each specimen and converted values into avian tetrahedral colorspace (u, s, m, l channels) using the vismodel function in pavo (Maia et al., 2013a) and based on a UV-sensitive visual system (Parrish et al., 1984). We finally converted quantum catches into perceptually uniform XYZ coordinates (Pike, 2012) for use in downstream comparative analyses (Figure 2A).

### Understanding tempo and mode of color evolution

To account for phylogenetic signal, we fit multivariate Brownian motion (BM) and Pagel’s λ models using the fit_t_pl function (Clavel et al., 2018). We compared models using generalized information criteria (GIC). The best-fitting model to the color data was a Pagel’s λ model (GIC weight ∼1.0; see Table S1). Therefore, we used this λ estimate to transform branch lengths of the phylogeny before running comparative analyses. To derive traits for downstream competition analyses and to visualize color pattern diversity, we used both ordinary principal components analysis (PCA) and phylogenetic PCA (pPCA). We performed pPCA using the phyl.pca_pl function in R (Clavel et al., 2018) and determined the number of principal components to retain by using a broken stick approach implemented in the vegan R package. For individual pPC traits, we fit univariate models using a combination of fitContinuous (Harmon et al., 2008), fit_t_comp (Drury et al., 2016), and OUwie (Beaulieu et al., 2012).

### Identifying drivers of color evolution

To understand the drivers of plumage color divergence among species, we used Bayesian phylogenetic mixed models (BPMMs) implemented in the MCMCglmm R package (Hadfield and Nakagawa, 2010). We used individual plumage patches as the unit of analysis. The multivariate response was the matrix of color XYZ coordinates, and the predictor variables were sex, color mechanism (structural versus pigment-based color), and insularity (mainland or island). We ran MCMC chains for 10^5^ generations and discarded the first 25% as burn-in. Convergence was checked visually using profile plots.

### Comparing plumage complexity with multivariate rates of color evolution

Using ancestral estimates of color divergence and plumage complexity, we used correlation tests to estimate the strength between them. To account for evolutionary variation in this relationship, we used a simulation-based approach similar to that used in the ratebystate function for univariate data (Revell, 2011). Briefly, we simulated evolution using the mvSIM function (Clavel et al., 2015) under Brownian motion 100 times based on the observed trait variance-covariance matrix estimated with phyl.vcv (Revell, 2011). For each simulation, we reconstructed ancestral states of coordinate-patch trait pairs, calculated rates and complexity as above, and created a null distribution for the correlation. To account for non-linearity of complexity metrics, we used Spearman’s *ρ* rather than Pearson’s r correlation coefficient. We determined P values as the proportion of null values greater than or equal to the observed value.

## Acknowledgements

We thank Ben Marks for assistance with bird specimens at the FMNH. We also thank Kristopher Menghi who helped with the collection of spectral data. This work was partially supported by grants from the National Science Foundation (NSF EP 2112468 to C.M.E. and S.J.H., NSF EP 2112467 and DEB 1557051 to M.J.A.).

## Data accessibility

All data sets and code for analyses are available on Dryad (DOI pending acceptance).

## Supporting Information

**Table S1.** Comparing models of color trait evolution. Generalize information criteria (GIC) scores were calculated using function in the RPANDA R package.

| Model | GIC | ΔGIC | GIC wt. | $\lambda$ |
| --- | --- | --- | --- | --- |
| Brownian motion | 13856 | 1385 | 0.00 | – |
| Pagel's $\lambda$ | 12472 | 0 | 1.00 | 0.16 |

**Table S2.** Rate-by-state correlation tests for alternative complexity metrics.

| **Metric** | **Spearman's $\rho$** | **P** |
| --- | --- | --- |
| Complexity spread-wing |  |  |
| Node-based | 0.37 | <0.01 |
| Tip-based | 0.58 | <0.01 |
| Complexity folded-wing |  |  |
| Node-based | 0.31 | <0.01 |
| Tip-based | 0.58 | <0.01 |
| Mean interpatch distance |  |  |
| Node-based | 0.39 | <0.01 |
| Tip-based | 0.58 | <0.01 |
| Mean color volume |  |  |
| Node-based | 0.34 | <0.01 |
| Tip-based | 0.54 | <0.01 |

**Table S3.** Phylogenetic linear regression results. Models fit with islands (island or continent distribution) as a predictor for different complexity metrics. See Methods for details.

| Plumage complexity metric | Estimate ± S.E. | P-value | $\lambda$ |
| --- | --- | --- | --- |
| Mean interpatch distance | -0.52 ± 0.44 | 0.24 | 0.55 |
| Mean color volume | -1.56 ± 3.75 | 0.68 | 0.29 |
| Color complexity (spread-wing) | 1.26 ± 0.85 | 0.14 | 0.50 |
| Color complexity (folded-wing) | 1.30 ± 0.92 | 0.16 | 0.49 |

**Figure S1.**
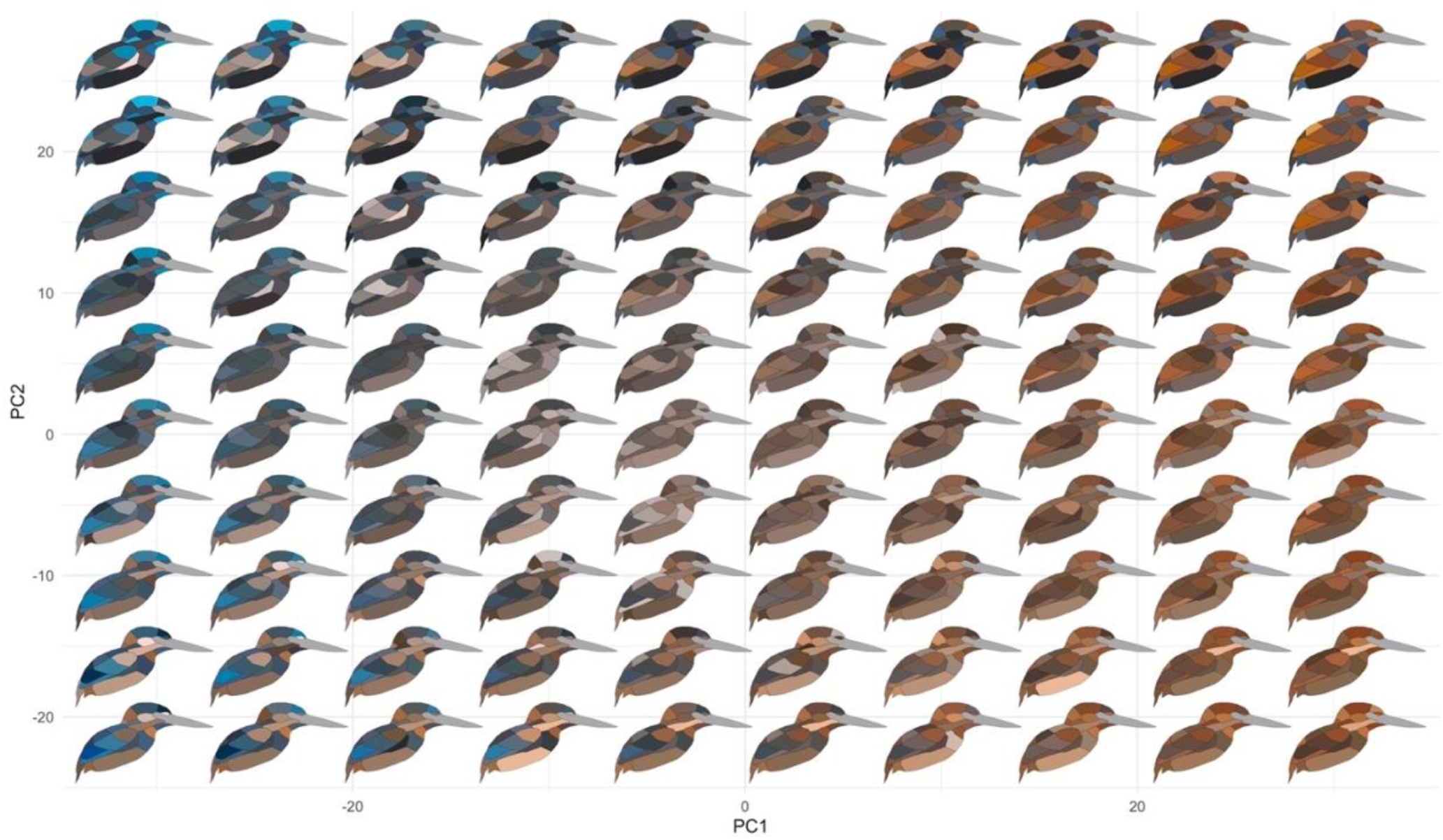
Color pattern morphospace of kingfishers. Bird images show depictions of color in a human visual system based on spectral measurements over a grid of ordinary principal components analysis (PCA) coordinates. Axes shown are PC axes 1 and 2, together accounting for >50% of plumage color variation in the clade.

**Figure S2.**
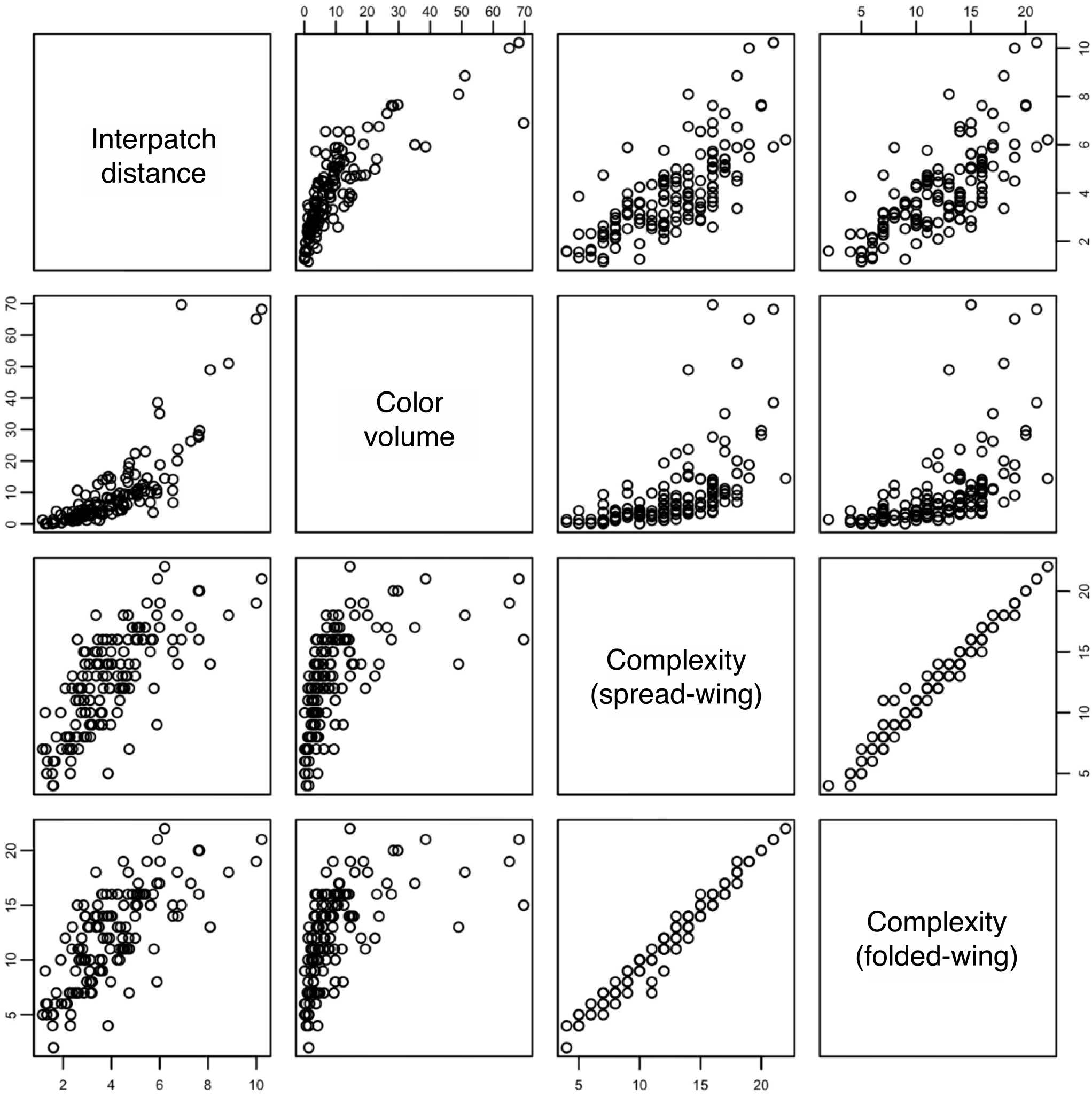
Pairwise scatter plots between plumage complexity metrics. Data shown for both tip-based (i.e. species) and node-based complexity scores along the phylogeny.

**Figure S3.**
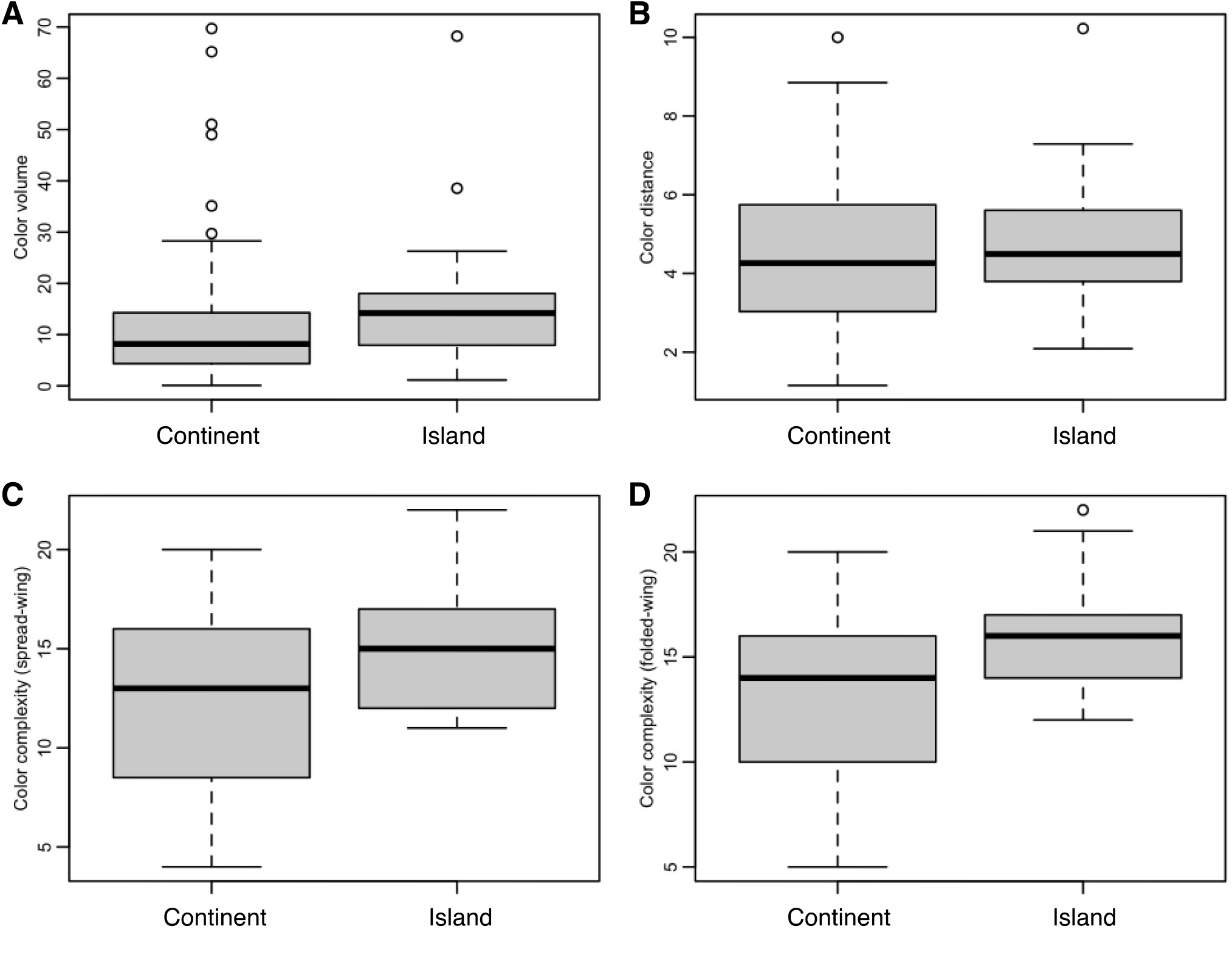
Islands and plumage complexity. There are no significant differences in plumage complexity between mainland and island species. Panels show different complexity metrics: plumage color volume (A), mean interpatch color distance (B), spread-wing plumage complexity (C), and folded-wing plumage complexity (D; see Methods for details).

**Figure S4.**
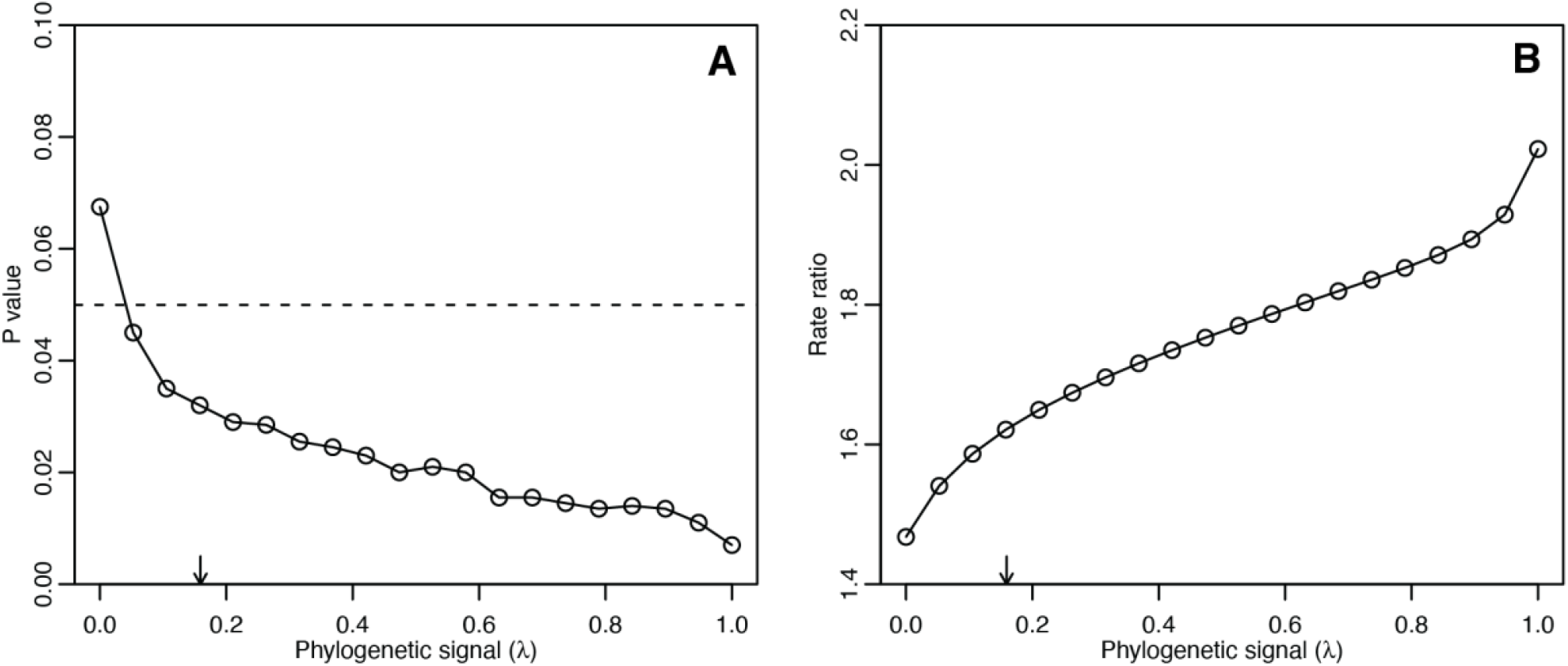
Effect of phylogenetic signal on rate tests. Panels show P values (left) and rate ratios (island vs. continents; right) as a function of phylogenetic signal. Arrows indicate empirical estimate of phylogenetic signal used in all comparative analyses.

**Figure S5.**
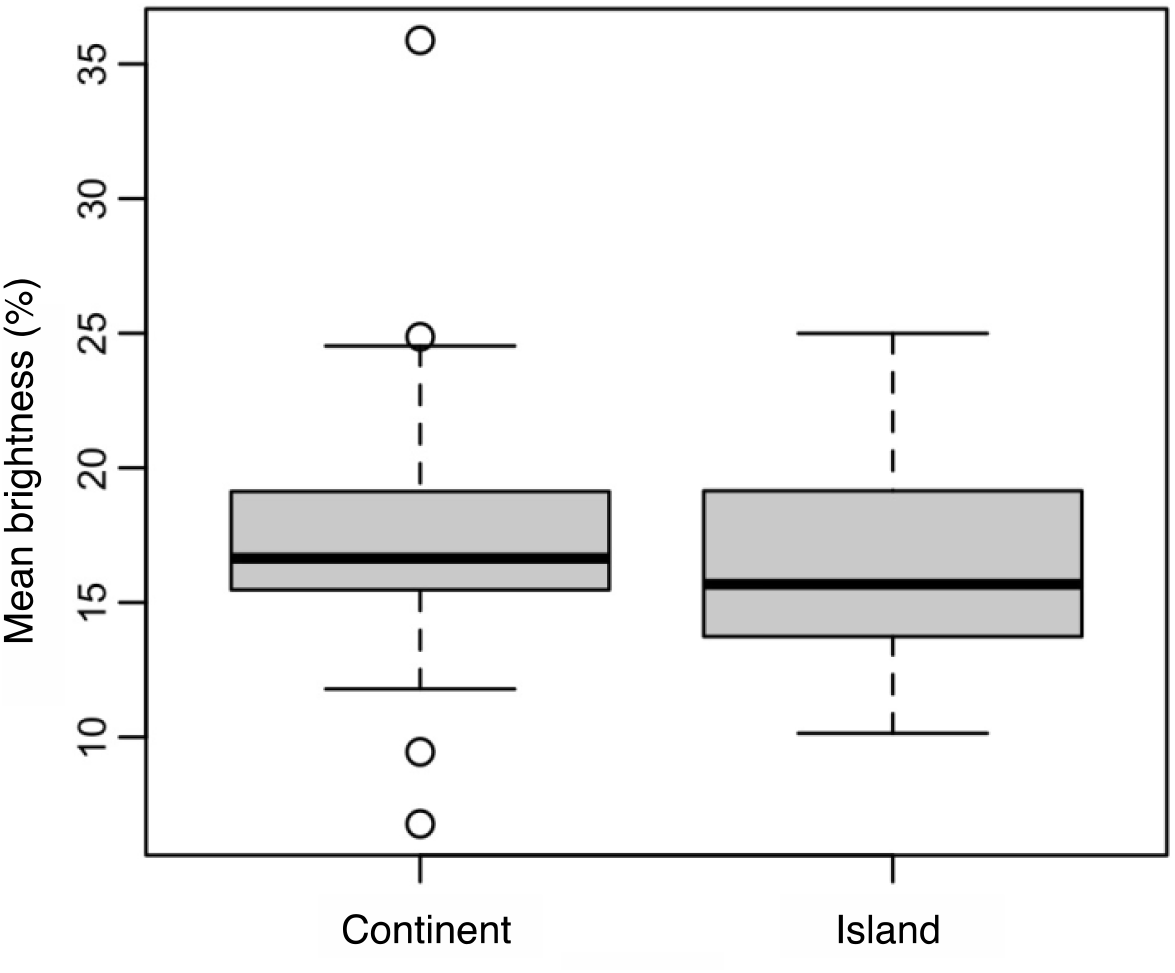
Species are equally bright on continents and islands. Phylogenetic linear model: estimate = -1.7, P = 0.11, λ = 0.50, σ^2^ = 0.78, N = 72.

**Figure S6.**
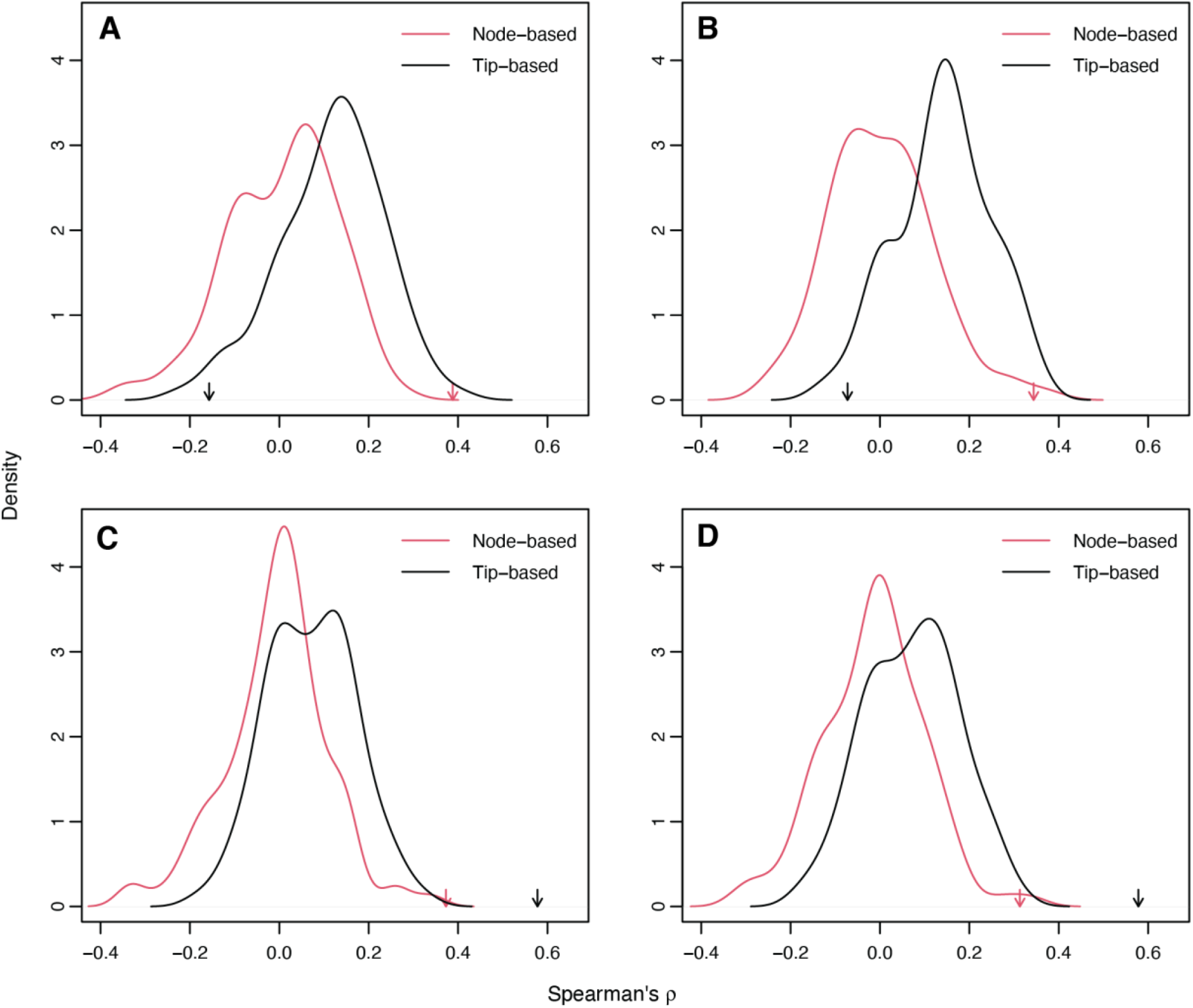
Effect of data analysis on rate-complexity correlations. Distributions show null correlation (Spearman’s ρ) inferred from simulated trait data for different complexity metrics: mean interpatch distance (A), mean color volume (B), spread-wing plumage complexity (C), and folded-wing plumage complexity (D). Line colors correspond to node-wise complexity scores inferred from single plumage patches (red lines) and ancestral complexity scores inferred from tip-only complexity scores (black lines). Arrows show observed correlation between rates and states.

